# Trends in animal model preference for preclinical drug testing for type-2 diabetes and future directions

**DOI:** 10.1101/2020.03.03.973230

**Authors:** Nuno Henrique Franco, Sónia Batista Miranda, Nora Kovacs, Attila Nagy, Folahanmi Tomiwa Akinsolu, I Anna S Olsson, Orsolya Varga

**Affiliations:** Department of Preventive Medicine, Faculty of Public Health, University of Debrecen, Hungary; i3S - Instituto de Investigação e Inovação em Saúde, Universidade do Porto; IBMC - Instituto de Biologia Molecular e Celular, Universidade do Porto

**Keywords:** Type-2 Diabetes Mellitus, animal models, animal research, preclinical testing, antidiabetic drugs, reproducibility, predictive validity

## Abstract

The importance of selecting accurate animal models of disease has become increasingly salient in recent years. This is particularly important for preclinical tests carried out to predict drug efficacy and safety for humans. We mapped past and current trends in animal model selection for type-2 Diabetes Mellitus (T2DM) antidiabetic drug research, and evaluate results in light of their implications for translational research.

From data gathered from published studies on 41 orally administered antidiabetic drug classes from the ‘MEDLINE Evaluator’ web service, we found animal model choice trends within – and across – drug classes to be substantially similar, with outbred rat strains being more prevalently used, since the 1990s. The observed consistency in choice of animal models in T2DM is advantageous for replicability and comparability of experiments, but prevalence of outbred strains raises a few concerns.

In face of current criticism of the translational value of animal tests, predictive validity must be assessed on a case-by-case basis. We address some of the issues pertaining the most popular animal models in antidiabetic drug development and testing, including the lack of information on their predictive validity, and propose a ‘back translation’ approach to estimate it retrospectively.

## Introduction

To understand the complex pathophysiological mechanisms of T2DM, as well as to develop new therapeutic approaches, scientists often resort to animal models reproducing different aspects of human diabetes, with varying fidelity. These models can either be spontaneous or induced, the latter by means of chemicals, highly-caloric diet, surgical manipulations (such as partial pancreatectomy) or genetic modification [1]. However, while there is a wide range of animal models available [2–4], it is less clear which are the most appropriate. The use of animal models for developing and testing – the potential efficacy, safety and appropriate dosing for humans – of therapeutic drugs is a mandatory requirement by regulators, to guarantee minimum safety standards for the licensing of drugs and chemicals. Such regulations include the FDA’s ‘Good Laboratory Practice for Nonclinical Laboratory Studies’ (21 CFR Part 58 [5]), and the EMA’s guidelines (ICH S6 (R1) [6]).

Reliable research on the efficacy and safety of candidate drugs warrants appropriate models. The FDA’s [7] Guidance for Industry on diabetes mellitus sets criteria on the phenotypic traits that appropriate animal models of T2DM should manifest, which include insulin resistance, hyperglycemia, and hyperinsulinemia. Recommended models include the leptin-deficient mouse (ob/ob), the leptin-receptor-deficient mouse (db/db), the obese Zucker rat (fa/fa), the Wistar Kyoto rat (fa/fa), and knockout mice lacking relevant targets, such as insulin receptors or glucose transporter 4 genes [7]. However, no specific ranking or method on how to assess the value of any of these models have so far been proposed. This significant gap might be closed by the Drug Development Tool (DDT) Qualification Programs, which was initiated in 2017. DDTs are measurements or methods (and associated materials) that aid drug development, including biomarkers, clinical outcome assessments, and animal models. For DDT considerations, an animal model is one in which a disease process/pathological condition can be investigated, and in which the disease/condition in multiple important aspects corresponds to the human disease/condition of interest. A qualified model can be recommended for efficacy testing in development programs for multiple investigational drugs for the same disease or condition. However, at this point, the qualification of an animal model through FDA’s Animal Model Qualification Program is voluntary, and the actual influence of the program has so far been marginal. To better understand trends in choice of animal models for drug therapy studies for type-2 diabetes (T2DM) mellitus, we gathered data on animal species and strains used in published studies on 41 orally administered antidiabetic drug classes, by a relevant MeSH term search on MEDLINE Evaluator data mining web service. We discuss these results in light of the known advantages and caveats of each type of model.

## Methods

We carried out an advanced search on the Meva (MEDLINE Evaluator) medico-scientific data mining web service. Meva, which does not act as MEDLINE portal, but as post-processor, allows querying PubMed with a specified search query, getting all abstracts, and generating frequency tables of MeSH terms [8]. MeSH is the National Library of Medicine’s controlled vocabulary thesaurus. MeSH descriptors are arranged in both an alphabetic and a hierarchical structure.

We selected 41 oral antidiabetic drug (OAD) from the seven main drug classes, namely: Sulfonylureas, Biguanides, Alpha glucosidase inhibitors, Thiazolidinediones, Dipeptidyl peptidase 4 (DPP-4) inhibitors, Glucagon-like peptide-1 (GLP-1) analogues, and Sodium-glucose co-transporter 2 (SGLT2) inhibitors.

The relevant MeSH term search query was as such: *[selected drug] AND “Diabetes Mellitus, Type 2”[Mesh] AND “Animals”[Mesh] AND “animals”[MeSHTerms:noexp] AND Journal Article[ptyp]*. Ranks based on absolute numbers of chosen animal models were classified in descending order. Proportions of animal models used were calculated with 95% confidence intervals for each strain/species. Cronbach’s alpha coefficient was calculated for each created OAD groups to explore the internal consistency of OADs, moreover an overall Cronbach’s alpha coefficient was calculated to check the inter-class consistency of OADs main groups. The actual ranking of animal models for each drug was compared with the raking of that drug class. Since the ranking for the overall of OADs was unknown, the average of rankings for drug classes was considered. Data on papers published until 2017 is presented.

## Results

The search criteria allowed retrieving articles to as back as the year 1965. The number of papers retrieved on animal models of T2DM followed an exponential growth over time (R^2^=0.958), although peaking in 2014 and dropping since then. Studies on therapeutic drugs on rodent models started rising in popularity from 1982, when mouse and rat studies together comprised 27%, until becoming the most popular models in 1988, and all years since then. Although mice were the most widely used rodent species until 1992, since then outbred rat strains have become the most popular models in T2DM drug tests, with the outbred Wistar and Sprague-Dawley strains being the most prevalent, followed by inbred wild-type C57BL/6 mice (Fig.1).

**Fig.1.**
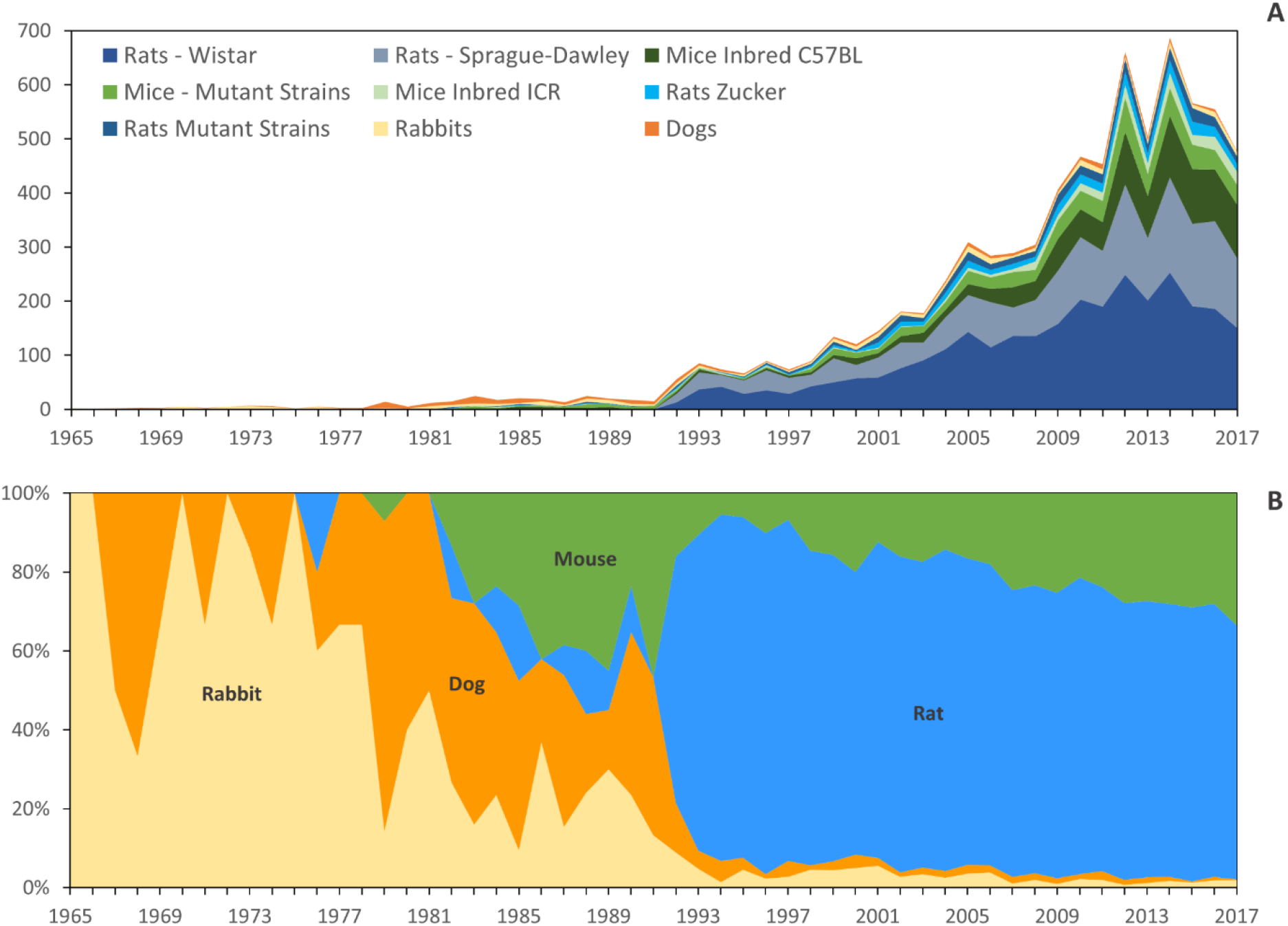
Number of publications for each species/strain, per year, for all drug classes. Above (Fig. 1A) is represented the absolute number of papers published on each animal model of T2DM, found in our search. Below (Fig 1B), global preferences for the main four species are represented, in terms of proportion of the overall number of studies.

The year 1992 marks not only a steep rise in number of papers on drug studies in animal models, but also a sharp decline in the use of non-rodent models, and particularly dogs and rabbits. Data on trends in choice of species provided as “Supplementary file 1”.

As regards consistency in choice of species/strain (of the 25 found) within drug classes, Wistar rats were the preferred strain for studies on three out of five drug classes (Table 1). We found a 0.91 ranking consistency between groups, indicating excellent internal consistency. At the level of each drug class, the lowest ranking consistency was found for alpha glucosidase inhibitors, with a coefficient of 0.59, while the highest was for thiazolidinediones, with 0.92. Detailed data on ranking is provided as “Supplementary File 2”.

**Table 1.** Ranking of animal models’ choice by antidiabetic groups. The table shows the five most frequently selected animal models (strains/species), in percentage, with confidence interval provided under the name of the strains/species. Cell colours indicate the species: blue-rat, green-mouse, yellow-rabbit, red-dog

|  | Rank |  |  |  |  |
| --- | --- | --- | --- | --- | --- |
|  | 1<sup>st</sup> | 2<sup>nd</sup> | 3<sup>rd</sup> | 4<sup>th</sup> | 5<sup>th</sup> |
| Sulfonylureas | Rats, Wistar<br>55.56 [50.49-60.63] | Rats, Sprague-Dawley<br>11.65 [8.38-14.93] | Rabbits<br>10.03 [6.96-13.09] | Mice, Inbred ICR<br>2.98 [1.25-4.72] | Dogs<br>2.71 [1.05-4.37] |
| Alpha glucosidase inhibitors | Rats, Wistar<br>39.51 [28.86-50.15] | Rats, Sprague-Dawley<br>17.28 [9.05-25.52] | Mice, Inbred C57BL<br>12.35 [5.18-19.51] | Mice, Inbred ICR<br>8.64 [2.52-14.76] | Rats, Mutant Strains<br>4.94 [0.22-9.66] |
| Thiazolidinediones | Mice, Inbred C57BL<br>16.26 [13-19.52] | Rats, Wistar<br>12.4 [9.49-15.31] | Rats, Mutant Strains<br>10.57 [7.85-13.29] | Rats, Zucker<br>9.76 [7.13-12.38] | Mice, Mutant Strains<br>9.55 [6.96-12.15] |
| DPP-4 inhibitors | Rats, Wistar<br>17.33 [12.39-22.28] | Mice, Inbred C57BL<br>12.44 [8.13-16.76] | Rats, Sprague-Dawley<br>11.11 [7-15.22] | Dogs<br>8 [4.46-11.54] | Mice, Inbred ICR<br>7.11 [3.75-10.47] |
|  |  |  |  |  | Rats, Mutant Strains<br>7.11 [3.75-10.47] |
|  |  |  |  |  | Rats, Zucker<br>7.11 [3.75-10.47] |
| GLP-1 analogues | Mice, Inbred C57BL<br>16.96 [12.98-20.94] | Rats, Sprague-Dawley<br>12.87 [9.32-16.41] | Rats, Mutant Strains<br>9.36 [6.27-12.44] | Rats, Zucker<br>9.06 [6.02-12.11] | Mice, Mutant Strains<br>8.19 [5.28-11.09] |

## Discussion

Contrary to the general trend in biomedical research, in which mice have mostly become the preferred species in the last 30 years, since the advent of the first transgenic mouse model in 1987 [9], rats are the most prevalently used animal species in T2DM drug development and testing. Our results also highlight the high-level of consistency for this choice of animal models. This consistency has the advantage of allowing direct comparison of different study results on the same drug, and thus of the reproducibility of said results, as well as facilitating meta-analyses. The inter-class consistency found is also useful when comparing either therapeutic or side effects between drugs of different classes. There are, however, important questions to address. While the pattern of selected rodent models fits the FDA’s [7] Guidance for Industry on diabetes mellitus, the prevalence of outbred strains should raise concern, as these are not genetically defined, and are more liable to genetic change than inbred strains, which impacts the reproducibility of results [10]. Also, due to their higher inter-individual variation, experiments on outbred strains are likely to have lower statistical power then similar sized experiments on inbred strains [10] thus lowering the probability of finding a true effect (therapeutic or toxicological) in tested drugs. It has, however, also been proposed that, at least for mouse models of diabetes, inbred and outbred mice appear to show comparable phenotypic variation, with variation being more influenced by strain choice and type of readout. [11]. If this is so, this should be factored in when choosing among the available models. Similar studies for rats – as well as confirmatory studies for available mouse strains – are, however, still warranted.

The advantages and limitations of each T2DM model for basic and applied research have been discussed in several review articles, most recently and thoroughly by Kleinert et al [12]. However, such reviews tend to focus on how closely animal models mimic the aetiology of T2DM (construct validity); or on how their phenotypes resemble human disease (face validity) [13]. Valuable as that information may be, the translational value of animal models should not be assumed to be a direct function of either construct or face validity (a view adequately dubbed as the ‘high-fidelity fallacy’ [14]), but assessed on a case-by-case basis. Hence, the criterion for selecting which animal models to use in preclinical studies should mainly be their predictive validity [15], i.e. the extent to which a given model can predict outcomes in the modelled system.

We hereby propose an approach for assessing the predictive validity of animal models of T2DM (applicable for other models of disease), by comparison of the same outcome measures in human and animal studies, an approach we have previously applied using Rosiglitazone as a case-study [16]. Briefly, we performed a search for both human and animal studies reporting rosiglitazone monotherapies with information on glucose and/or HbA1c values, which returned 71 results. Information from all included studies was extracted and entered into a single database with as many observations for each study as the number of fasting glucose and/or HbA1c outcome measurements reported at distinct follow-up times in that study. Linear regression models were fitted on human observations for both outcomes and predictive value of animal models was assessed by comparing the same outcome measures from human trials and animal studies. Findings showed that although the consistency of animal species-based models with the human reference for glucose and HbA1c treatment effects is highly variable, glucose and HbA1c treatment effects in rats agreed better with the expected values based on human data than in other species. In addition, rats showed significantly lower scores of deviation from the human reference than mice for glucose and HbA1c treatment effects. The question regarding which strain is the most appropriate to model the clinical efficacy of rosiglitazone, however, could only be tentatively answered. There was no statistical difference in deviation scores observed between rat strains. Among mouse strains, C57BL/6 showed the most consistent, while db/db showed the least consistent results.

This insight provided by our comparative approach, allows us to suggest that the prevalence in the use of rats – including over larger and phylogenetically closer mammal species – may be adequate for preclinical drug testing, at least for some drug classes and outcomes. However, it also highlights the need to critically assess their predictive value for different outcomes and drug classes, on a case-by-case basis. This ‘back translation’ (i.e. from clinical to preclinical trials) approach has been proposed for retrospectively identifying possible causes of translational failure, as well as to further refine successfully predictive models [15]. More importantly, we propose it can be used to facilitate an informed choice of appropriate animal models, based on their predictive validity for different research aims, types of T2DM manifestations and aetiologies, and pharmacological principles, among other parameters. Such detailed information could be organised in a comprehensive database of the predictive validity of current models, animal or otherwise, to improve the translational value and overall quality of preclinical research.

